# Hypermut 3: Identifying specific mutational patterns in a defined nucleotide context that allows multistate characters

**DOI:** 10.1101/2024.10.24.620069

**Authors:** Zena Lapp, Hyejin Yoon, Brian Foley, Thomas Leitner

## Abstract

**Summary:** The detection of APOBEC3F- and APOBEC3G-induced mutations in virus sequences is useful for identifying hypermutated sequences. These sequences are not representative of viral evolution and can therefore alter the results of downstream sequence analyses if included. We previously published the software Hypermut, which detects hypermutation events in sequences relative to a reference. Two versions of this method are available as a webtool. Neither of these methods consider multistate characters or gaps in the sequence alignment. Here, we present an updated, user-friendly web and command-line version of Hypermut with functionality to handle multistate characters and gaps in the sequence alignment. This tool allows for straightforward integration of hypermutation detection into sequence analysis pipelines. As with the previous tool, while the main purpose is to identify G to A hypermutation events, any mutational pattern and context can be specified.

**Availability and implementation:** Hypermut 3 is written in Python 3. It is available as a command-line tool at https://github.com/MolEvolEpid/hypermut3 and as a webtool at https://www.hiv.lanl.gov/content/sequence/HYPERMUT/hypermutv3.html.

## Introduction

As part of the human immune response, APOBEC3F and APOBEC3G proteins can deaminate cytosine residues to uracil on viral DNA, leading to a guanine to adenine mutation in subsequent viral sequences (Refsland *et al*., 2012). These mutations usually occur in the context of downstream RD (A or G, not C) nucleotides (Refsland *et al*., 2012; Yu *et al*., 2004). This pattern of hypermutation has been detected in sequences from human immunodeficiency virus (HIV) (Vartanian *et al*., 1991), hepatitis B virus (Noguchi *et al*., 2005), and mpox (Desingu *et al*., 2024), among others. Since these mutations do not arise from standard vertical evolution of viral mutations during replication, they need to be removed or modified prior to performing analyses that use mutations to estimate relatedness between sequences, such as computing phylogenies, identifying transmission clusters, or dating latent proviral sequences.

We previously published a webtool, Hypermut (Rose and Korber, 2000), that detects patterns consistent with hypermutation in genome sequences. Given an alignment including a *reference sequence* that is assumed to have no signature of hypermutation, Hypermut detects potentially hypermutated positions in each *query sequence* in the alignment. A decade later, we updated the webtool (unpublished) to enable users to compare the number of mutations occurring in user-defined upstream and downstream nucleotide *primary contexts* vs. *control contexts* (**Figure 1A**). The context can be enforced in the reference sequence, the query sequence, or both. The use of multistate characters in the user-defined contexts indicate that any of the included nucleotides may be considered a match in the alignment. If the correct mutation occurs in a primary context (**Figure 1B**, q1), it is considered a primary match. If an incorrect mutation, or no mutation, occurs in a primary context (**Figure 1B**, q2), it is considered a potential primary match (but not a match). The same logic is followed to identify potential and actual matches for control contexts (**Figure 1B**, q3). Hypermut 2 returns a summary of the number of mutations in contexts of interest compared to control contexts for each sequence and uses these numbers to compute Fisher’s exact p-values that quantify whether a sequence has more mutations than expected in contexts of interest compared to control contexts. Users can also download a file including the positions of potential mutation sites in the primary and control contexts, and whether there was a mutation match at the site. However, Hypermut 2 only matches to ACGT characters in the alignment and therefore does not consider sites with multistate characters or gaps, which may occur due to virus population diversity.

**Figure 1:**
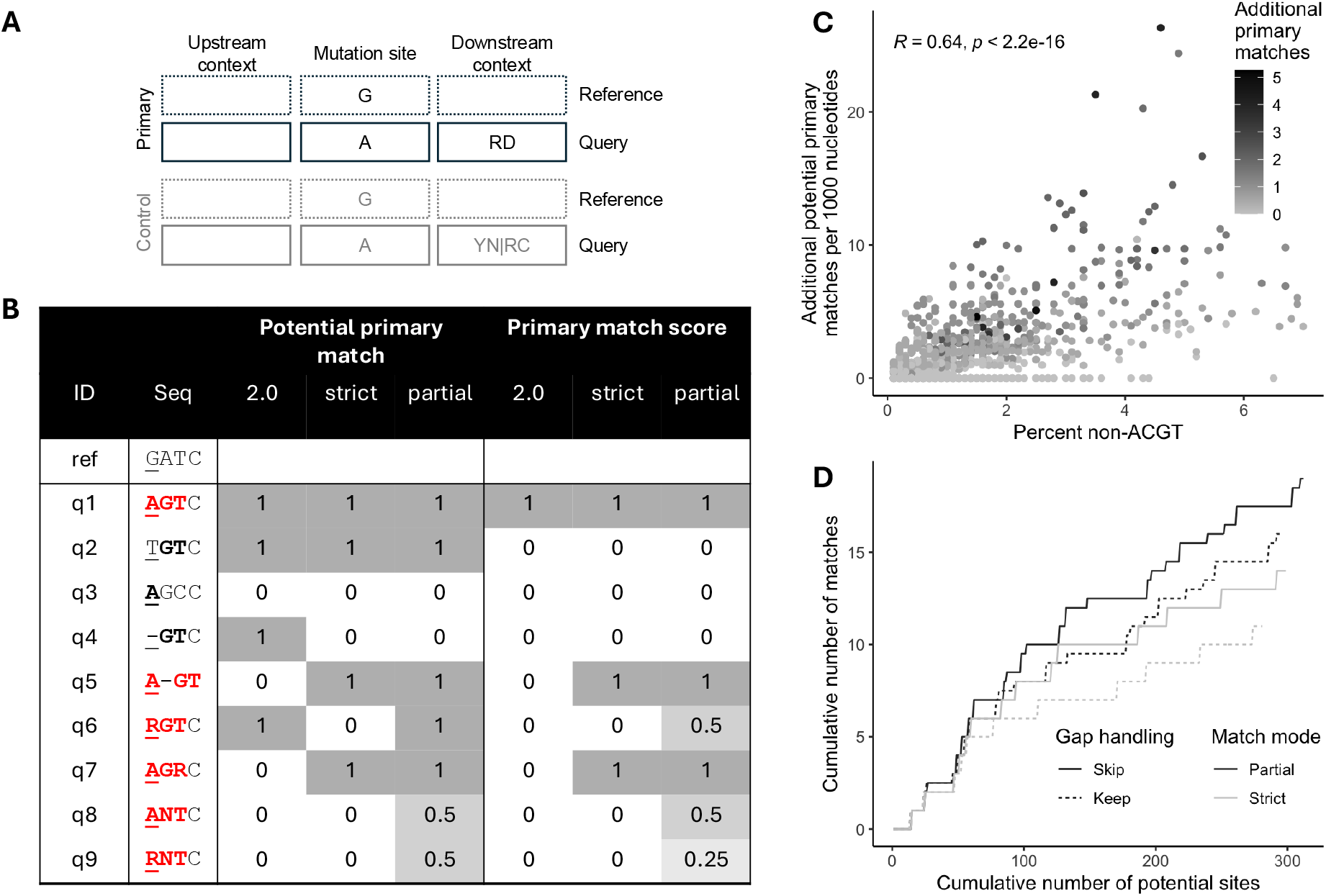
Hypermut 3 overview. **(A)** Example mutation site and context (for APOBEC3G and APOBEC3F) using IUPAC codes. The control pattern is the complement of the primary pattern and inferred by the program. **(B)** Example reference and query sequences, and whether they are considered a potential primary match and an actual primary match for three Hypermut versions: 2.0, 3.0 strict, and 3.0 partial. These numbers assume that gaps are skipped (for Hypermut 3) and that the context is enforced on the query sequence. In the sequence, the underlined nucleotide is the mutation site, correct mutations or contexts are bolded, and matches for Hypermut 3 partial mode are colored in red. The IUPAC code R indicates A and/or G, and the IUPAC code N indicates any of the bases. **(C)** Correlation between percent non-ACGT characters in the query sequence and the number of additional potential primary matches identified when using partial matching compared to strict matching. The color indicates the number of additional primary matches observed. **(D)** For an example sequence, the cumulative number of potential sites vs. the cumulative number of actual matches identified for each combination of partial vs. strict matching (color) and keeping vs. skipping gaps in the alignment (line type).

Since the original publication of Hypermut 25 years ago, there has been an explosion in available sequencing data and corresponding development of automated bioinformatic pipelines. Here, we present a substantial update to Hypermut that allows for the integration of hypermutation detection into automated bioinformatic pipelines, and that can handle multistate characters and alignment gaps.

## New developments in Hypermut 3

Hypermut 3 extends Hypermut 2 by optionally handling gaps and multistate characters in the alignment. Furthermore, it is available as a command-line tool that can be straightforwardly integrated into bioinformatic pipelines. We have simplified the input compared to Hypermut 2 by automatically setting the control context to be the exact complement of the primary context, which will reduce accidental user error for complex control patterns. The output information remains the same.

### Gap handling

Gaps in the mutation site are not considered (**Figure 1B**, q4). By default, gaps in the context are ignored when considering potential mutation sites of interest (**Figure 1B**, q5), i.e., gaps are skipped over and the following characters in the sequences are used for the context patterns. We also provide an option to keep gaps, using them as characters, in which case gaps in the context are not ignored.

### Multistate character handling

Matching at positions with multistate characters in the alignment can be handled in two ways (**Figure 1B**, q6-q9): (1) *strict mode* ignores positions where the multistate character in the sequence is broader than the user-defined pattern, and (2) *partial mode* identifies partial matches for positions with some overlap between the multistate character and the user-defined pattern. Only strict matching can be performed if the reference sequence contains multistate characters.

#### Strict mode

In strict mode, sites with multistate characters are considered only when the nucleotides that make up the multistate character are entirely included in the user-defined search pattern. In cases where the multistate characters contain nucleotides that are not included in the user-defined search pattern, the location is ignored and not considered a potential primary or control match.

#### Partial mode

Partial mode returns the same complete matches as strict mode, but also assigns partial matches to locations with multistate characters that contain some, but not all, nucleotides in the user-defined search pattern. For this mode, we assume that each nucleotide is present at equal frequency in the population, and that all potential combinations of nucleotides are also present at equal frequency. In this case, there may be partial matching in the context or the mutation site. Partial matching in the context leads to a fractional *potential match* “count,” while partial matching in the mutation site leads to a fractional *actual match* “count.” For a given position, we quantify the extent of matching by determining the proportion *P*_*s,c*_ of standard nucleotides (ACGT) in the IUPAC code of the sequence that are in the correct context:

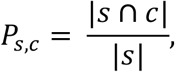

where *s* is the set of nucleotides present in the IUPAC code for the sequence, *c* is the set of nucleotides present in the IUPAC code for the context or mutation of interest, and |·| is the cardinality (i.e., length) of the set. For a given context of length *l*, the potential match “count” *M* is the product of the proportions for each position in the context:

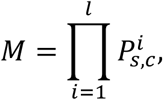

where 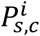 is the proportion of matching nucleotides for the *i*^th^ position in the context pattern. If there are multiple possible contexts, then the total potential match count is the sum of the individual match counts. The actual match count is computed as:

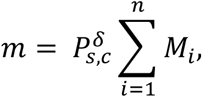

where δ designates the mutation of interest, *n* is the number of possible contexts, and *M*_*i*_ is the potential match count for context *i*.

Potential and actual match counts are computed for the user-defined primary context as well as the inferred control context.

## Availability and implementation

Hypermut 3 is written in Python 3 and requires the scipy package (Virtanen *et al*., 2020). It is available as a command-line tool at https://github.com/MolEvolEpid/hypermut3 and as a webtool at https://www.hiv.lanl.gov/content/sequence/HYPERMUT/hypermutv3.html.

## Example of use

To investigate the effects of gaps and multistate characters on the detection of potentially hypermutated positions, we downloaded from the LANL HIV database 1541 high-quality HIV-1 *gag* and *env* population sequences aligned to HXB2 (GenBank accession number K03455; all data available at https://github.com/MolEvolEpid/hypermut3/tree/main/manuscript/data). Due to the diversity of the sequences in the alignment, the mean percent of gaps within a query sequence was 24.9% (range: 0%-40%). The mean percent of non-ACGT characters was 1.1% (range: 0.06%-7.0%). We ran Hypermut 3 on these sequences, using HXB2 as the reference sequence, in both strict and partial modes, with and without skipping gaps in the context.

We first investigated whether sequences had signatures of hypermutation using a Fisher’s exact p-value threshold of 0.05. When keeping gaps in the context, 2 sequences fell below the threshold in both strict and partial modes, and 1 additional sequence fell below the threshold in partial mode only. When skipping gaps in the context, 8 sequences had signatures of hypermutation in both strict and partial modes.

In general, skipping gaps increased the number of potential contexts in 94.7% (1460/1541) of sequences in both strict and partial modes, and the number of actual matches in over 70% of sequences (strict: 1132/1541, 73.5%; partial: 1154/1541, 74.9%).

We next compared potential and actual matches for the strict and partial modes, both with gap skipping. When compared to the strict mode, using the partial mode led to 49.2% (758/1541) of sequences having more potential contexts and 40.2% (619/1541) of sequences having more potential hypermutations. As expected, we observed a positive correlation between the percent of non-ACGT characters and the number of additional potential sites (**Figure 1C**). An example of the difference in the cumulative number of potential sites and matches under different multistate match mode and gap handling conditions is shown in **Figure 1D**.

### Potential applications

Hypermut 3 can be used to detect APOBEC3F- and APOBEC3G-induced mutations as well as other mutations of interest. Among other applications, the output from Hypermut may be useful as a quality control step in a sequence analysis pipeline to remove sequences with certain mutational signatures or to mask certain positions within sequences, thus avoiding biases in downstream results. The new partial matching mode is particularly useful for identifying potential hypermutation in viral population sequences derived from, for example, Sanger sequencing of a virus population or a resolved consensus sequence of Illumina reads.

The web version of Hypermut 3 is straightforward to use for even those with no programming background and provides a means to easily identify potentially hypermutated sequences from an alignment of interest. For those with more programming experience, we also provide a command-line version of Hypermut 3 that can be incorporated into bioinformatic pipelines. This greatly expands the potential applications of Hypermut by allowing it to become a standard tool used to automatically identify and flag hypermutated sequences in real time for purposes ranging from research to public health.

## Conclusion

We have updated the original Hypermut software to accommodate the increase in genome sequencing and the development of automated bioinformatic pipelines. The added functionality of Hypermut 3 compared to the original published version allows for the identification of potential hypermutation events for multistate characters (as of Hypermut 3), provides a comparison of mutations in primary and control contexts to more easily identify hypermutated sequences (as of Hypermut 2), and enables straightforward integration into bioinformatic pipelines (as of Hypermut 3). Positions or sequences identified as likely hypermutants can then be removed or transformed prior to further sequence analyses, thus reducing potential biases in the downstream results.

## Funding

This work was supported by the National Institutes of Health [grant R01AI087520 to TL, interagency agreement AAI24007-001-00000 HIV/SIV, Database and Analysis Unit to HY and BF] and the Los Alamos National Laboratory [Laboratory Directed Research and Development program fellowship project no. 20230873PRD4 to ZL].

## Notes

https://github.com/MolEvolEpid/hypermut3/

https://www.hiv.lanl.gov/content/sequence/HYPERMUT/hypermutv3.html

### Competing Interest Statement

The authors have declared no competing interest.

